# Potential impacts of transport infrastructure and traffic on bird conservation in Swedish Special Protection Areas

**DOI:** 10.1101/351692

**Authors:** Jan Olof Helldin

## Abstract

The ecological impacts of roads and railways extend into the surrounding landscape, leading to habitat degradation and reduced animal densities within an area that is considerably larger than the actual road or railway corridor. For birds, an extensive meta-analysis has pointed at an average of 20% density reduction within 1 km from the infrastructure. I investigated to what extent this density reduction could potentially compromise the habitat quality and conservation value of Swedish Natura 2000 areas designated for the protection of birds (Special Protection Areas; SPAs). A majority (63%) of Swedish SPAs are at least to some extent found within this 1 km potential effect zone. The total overlap between SPA and effect zone is 126,000 ha, or 4.2% of the country’s SPA area. There are however large differences among biogeographical regions. In the southern (continental) and coastal regions combined, 25.8% of the total SPA area fall within the effect zone, representing an estimated 4-7% reduction in bird abundance within SPAs. The probability of overlap with effect zone is higher for larger SPAs. However, the proportion of overlap is higher for smaller SPAs, and accordingly smaller sites can be assumed to experience a greater impact from transport infrastructure and traffic. The impacts on Natura 2000 sites are particularly concerning as this network of protected areas is a cornerstone to maintain and restore biodiversity within EU. I recommend putting a stronger emphasis in the management of Natura 2000 sites on the potential threats to wildlife conservation caused by transport infrastructure and traffic. Special attention should be paid in sites with a large overlap with the effect zone, and in sites hosting particularly vulnerable taxa or habitats. Infrastructure owners and managers should do their best to minimize and compensate for the negative impacts of roads and railways and related traffic in SPAs and other protected areas.

## Introduction

### Ecological impact of transport infrastructures

Infrastructure development is recognized as one of the significant drivers of the global biodiversity loss, and with increasing traffic and expanding infrastructure networks worldwide the pressure on biodiversity is expected to increase in the nearest decades (EEA 2011, 2012, OECD 2012). The impacts of transport infrastructures on wildlife are well described (reviewed by, e.g., Forman et al. 2003, van der Ree et al. 2015), and include loss of habitat, traffic casualties, creation of physical barriers, disturbance by noise, light and other visual cues, spread of chemicals, dust and alien species, changes in hydrology and microclimate, and accidental spill. Most of these impacts extend into the surrounding landscape, leading to a degradation and fragmentation of habitats, and for some animal species to restricted movements, increased mortality, and avoidance of a zone around the infrastructure (Forman et al. 2003, EEA 2011, van der Ree et al. 2015).

Due to these impacts, the population densities of many animal species are reduced within a distance from larger infrastructures (Rytwinski & Fahrig 2015). For example, the population densities may be reduced on up to about 1 km distance for birds (Forman & Deblinger 2000, Forman et al. 2002, Benítez-López et al. 2010) and anurans (Eigenbrod et al. 2009), and up to 5 km for mammals (Benítez-López et al. 2010). Not only large infrastructures but also minor and unpaved roads may have a considerable impact on some wildlife species (e.g., van Langevelde & Jaarsma 2009, Benítez-López et al. 2010, Shanley & Pyare 2011). Based on an extensive meta-analysis, Benítez-López et al. (2010) showed that the mean bird and mammal abundance in an effect zone around infrastructure is reduced by 20-30%, and with an increasing reduction with proximity to the infrastructure (see Fig. 1).

**Figure 1.**
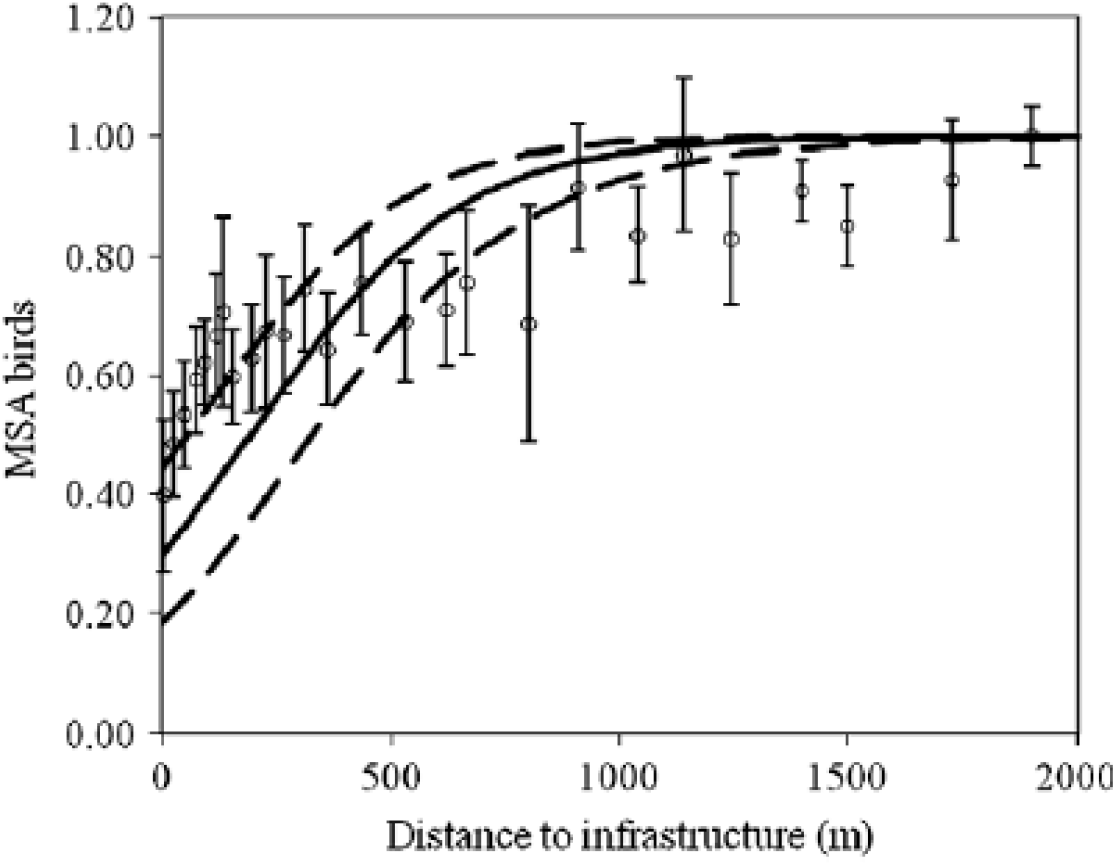
Mean species abundance (MSA) of birds as a function of distance to infrastructure (logistic regression). Open dots represent the pooled results of a meta-analysis per distance interval ±SE. Solid black line denotes the estimated curve for the decline of MSA in proximity to infrastructure; dashed lines are the 95% upper and lower limits of the confidence bands of the curve. Figure from Benítez-López et al. (2010).

Accordingly, in regions with dense infrastructure networks, large natural areas may be situated within this effect zone and therefore impoverished in species sensitive to traffic and transport infrastructures. For example, in the United States, the road effect zone covers 15-22% of the total land area, and more than 60% of some particularly exposed biomes, such as coastal regions and river basins (Forman 2000, Riiters & Wickham 2003). In Spain, a country with intermediate road density with European standards, reduced bird densities due to transportation infrastructure are expected in 49% of the country, and reduced mammal densities in as much as 96% of the country (Torres et al. 2016). Some habitats of particular importance to biodiversity in Europe, such as wetlands, semi-natural grasslands and temperate broad-leaved forest, may be disproportionately affected by roads because of the landscape structure (Helldin et al. 2013, Karlson & Mörtberg 2015, Torres et al. 2016). Disturbance (noise and visual cues) tend to be stronger and extend further from the road in open habitats as compared to forest (Forman & Deblinger 2000, Reijnen & Foppen 2006). There is a growing concern globally about impact of roads and traffic on wildlife populations in protected areas and other biodiversity hotspots (Forman & Deblinger 2000, Ament et al. 2008, Selva et al. 2011, Laurence & Balmford 2013, Bager et al. 2015, Gadd 2015, Seshadri & Ganesh 2015).

### Integrity of the Natura 2000 network

The Natura 2000 network of protected areas is a key tool in the maintenance and restoration of biodiversity within the European Union (EU). The network consists of Special Protection Areas (SPAs) designated according to the Birds Directive, and Special Areas of Conservation (SACs) designated according to the Habitats Directive (EEC 1992, 2010). Under these directives, the network is supposed to provide protection for vulnerable wildlife and habitats. One important motivation for designating Natura 2000 sites, particularly in coastal and other lower-elevation areas, is the protection against negative impacts of urbanization and infrastructure development (EEA 2012). There are presently more than 26,400 sites in the Natura 2000 network, accounting for *ca* 18 % of EUs land territory.

The network is however biased toward highland areas, and lowland areas are underrepresented (Oldfield et al. 2004, Mariorano et al. 2007). In addition, the average size of the Natura 2000 sites is quite low, and particularly so in lowlands (Maiorano et al. 2007, Gaston et al. 2008, EEA 2012). As smaller sites are more susceptible to pressure from land use and human activities surrounding them, major concerns are expressed about the capacity of existing protected areas to maintain their biodiversity values (Chape et al. 2005, Gaston et al. 2008, Maiorano et al. 2008, Kati et al. 2015).

Though many assessments of the Natura 2000 network’s effectiveness in protecting species have been reported in the last years, a vast majority of these relied on rather static population data, such as species’ geographic distribution, species presence/absence, or habitat suitability (e.g., Araujo et al. 2007, Maiorano et al. 2007, 2015, Sánchez-Fernández et al. 2008, Iojă et al 2010, López-López et al. 2011, Gruber et al. 2012, Albuquerque et al. 2013, D’Amen et al. 2013, Lison et al. 2013, Rubio-Salcedo et al. 2013, Trochet & Schmeller 2013), and accordingly were designed to assess ecological representativeness of the network rather than tracking changes in population densities due to environmental impacts. In view of the many negative population trajectories reported for both avian and non-avian species in the EU (EEA 2015), it appears necessary to analyze the ecological functionality of the Natura 2000 network with regard to pressures both within and outside the designated areas, but few studies have taken this course.

Frequency of transport infrastructure within Natura 2000 sites was investigated by Tsiafouli et al. (2013), showing that as a European average, roads are present in 29% of the sites, with a higher frequency in the countries in the south. A preceding study of Greek Natura 2000 sites, by Votsi et al. (2012), showed that 85% of sites are bisected by roads. Insufficient functional connectivity of the Natura 2000 network has been reported by Gurrutxaga et al. (2011) and Opermanis et al. (2012), suggesting that dispersal barriers exist between sites, for example in the form of large roads (Gurrutxaga et al. 2011). With regard to disturbance, the European Environmental Agency estimated that almost 20% of Natura 2000 areas are presently adversely affected by high levels of environmental noise, largely owing to major transport infrastructures (EEA 2016). In an assessment of the impact on Natura 2000 sites of major traffic arteries planned or under construction as part of EU’s TEN-T framework, Byron & Arnold (2008) estimated that 379 SPAs (8% of sites) and 953 SCIs (4% of sites) would be adversely affected by these new traffic arteries, with potential effects also on Natura 2000 network coherence.

### Aim of the study

Sweden is one of the European countries that are least fragmented by transport infrastructures and built-up areas (EEA 2011), and the physical impact of infrastructure is generally not well acknowledged in Swedish nature conservation. It is not until recently that national status reports for biodiversity have described the threats from infrastructure and traffic on species conservation (Bernes 2011, Naturvårdsverket 2015a), and the current national conservation action plan contains few requirements to infrastructure managers (Naturvårdsverket 2015b). Biodiversity is generally insufficiently described in impact assessments of transportation infrastructure plans and projects (Wärnbeck 2013, Karlson et al. 2014). Few, if any, management plans for protected areas address the full array of potential ecological impacts of transport infrastructures on the areas’ conservation status or management (Helldin & Tytor 2017). This ignorance is not unique for Sweden, but appear to be largely similar in most EU member states (Tsiafouli et al. 2013, EEA 2015; but see, e.g., Selva et al. 2011, Votsi et al. 2012).

In order to illustrate and highlight the impacts of infrastructure on protected areas in particular, I estimated the frequency and proportion of Swedish SPAs situated within the potential effect zone for birds around existing larger transport infrastructures (roads and railways), and hence can be expected not to reach their full conservation potential due to infrastructure impacts. I included only SPAs in the estimation, i.e., areas designated specifically for the protection of birds, because the effects of roads and railways on birds are well described in literature and apparently can impact a majority of bird species (Reijnen & Foppen 2006, Benítez-López et al. 2010, Rytwinski & Fahrig 2015). I assessed the conservation value of SPAs that is lost due to infrastructure in terms of reduced predicted bird abundance. Because of the large geographic variation over the country in density of the infrastructure network and proportion of area within SPA, I separated the analyses between biogeographical regions. I tentatively explored the association between the infrastructure impact on an individual SPA and its size and dominating habitats. In this paper, I present the results of these analyses and propose improvements for the management of Swedish SPAs.

## Methods

A map of the effect zone for birds around the existing larger Swedish transport infrastructure was produced. I assumed a standardized effect distance of 1 km from the infrastructure, following the result from a meta-analysis presented by Benítez-López et al. (2010). To my knowledge, this is the most comprehensive analysis of infrastructure impacts on bird densities, including 49 bird datasets and 201 bird species. Most studies in the meta-analysis were conducted in biomes that occur in Sweden, i.e., taiga, temperate broadleaf forest or alpine/tundra (39 of the 49 datasets), and on species that occur in Sweden (105 of the 201 species), and I therefore judged the results relevant in a Swedish perspective. The results have also previously been applied to assess the impacts of the road network on birds in Sweden (Karlson & Mörtberg 2015) and Europe (Torres et al. 2016). I consider the assumption of a 1 km effect distance to be conservative because i) individual studies in the meta-analysis indicated reduced bird populations on larger distances, and ii) impacts on longer distances may not necessarily result in direct population declines, but yet be of ecological significance. Infrastructure data were obtained from Open Street Map (http://openstreetmapdata.com), using only the following road classes: primary road, secondary road, tertiary road, motorway, trunk road, railway (thus excluding minor roads for which ecological effects are less well known).

The effect zone map was overlaid with all Swedish SPAs to calculate the area and proportion of each site situated within the effect zone. Shape of areas and habitat distribution were obtained from European Environment Agency’s Natura 2000 database (http://www.eea.europa.eu/data-and-maps/data/natura-2000-eunis-database). SPAs were separated depending on biome (based on a combination of the Natura 2000 database and Global Biomes data from the CIESIN; http://sedac.ciesin.columbia.edu/data/set/nagdc-population-landscape-climate-estimates-v3/maps?facets=theme:climate) and proximity to coast (data on coastline obtained from Open Street Map; http://openstreetmapdata.com) on the following terms:

- *continental region:* sites with >50% of the area within EU continental region,
- *mixed-forest region:* sites with >50% of the area within EU boreal region and in CIESIN broadleaf and mixed-forest region,
- *boreal region:* rest of sites within EU boreal region but with no part within EU alpine region, or
- *alpine region:* sites with at least some part of the area within EU alpine region; in combination with
- *coastal:* mainland sites with at least some part within 20 km from coast of mainland Sweden (including mainland islands Öland and Gotland), or
- *inland:* the rest of mainland sites.

The alpine region in Sweden is only inland, i.e., no alpine coastal sites exist. Because only two continental sites are inland, all continental sites were pooled in one region. In addition, an off-coastal region was formed including all sites with no contact with mainland Sweden irrespective of terrestrial biome. Hereby a total of seven biogeographical regions were obtained (see Table 1 and Fig. 2).

**Table 1.** Number, area and proportion of Swedish SPAs within an assumed effect zone of 1 km from larger transport infrastructure, and predicted total reduction in bird abundance (with 95% confidence interval) due to the effects. Results are given for the entire country and separated by biogeographical region.

| Region | Total<br>no. of<br>SPAs | Total area<br>of SPA<br>(ha) | No. of SPAs<br>affected to |  | Total area<br>of SPA in<br>effect zone<br>(ha) | Proportion<br>of total SPA<br>in effect<br>zone (%) | Reduction in<br>bird<br>abundance<br>(% with C.I.) |
| --- | --- | --- | --- | --- | --- | --- | --- |
|  |  |  | >0% | >50% |  |  |  |
| Continental | 41 | 53,331 | 41 | 18 | 19,202 | 36.0 | 7.2 (4.3-11.9) |
| Mixed-forest coastal | 94 | 103,064 | 71 | 24 | 20,865 | 20.2 | 4.0 (2.4-6.7) |
| Mixed-forest inland | 161 | 258,804 | 119 | 43 | 34,622 | 13.4 | 2.7 (1.6-4.4) |
| Boreal coastal | 22 | 21,712 | 15 | 8 | 5,968 | 27.5 | 5.5 (3.3-9.1) |
| Boreal inland | 130 | 162,924 | 63 | 30 | 13,122 | 8.1 | 1.6 (1.0-2.7) |
| Alpine (only inland) | 26 | 1,984,005 | 16 | 0 | 28,242 | 1.4 | 0.3 (0.2-0.5) |
| Off-coastal | 64 | 415,308 | 14 | 0 | 3,913 | 1.0 | 0.2 (0.1-0.3) |
| *All Sweden* | *538* | *2,999,149* | *339* | *123* | *125,946* | *4.2* | *0.8 (0.5-1.4)* |

**Figure 2.**
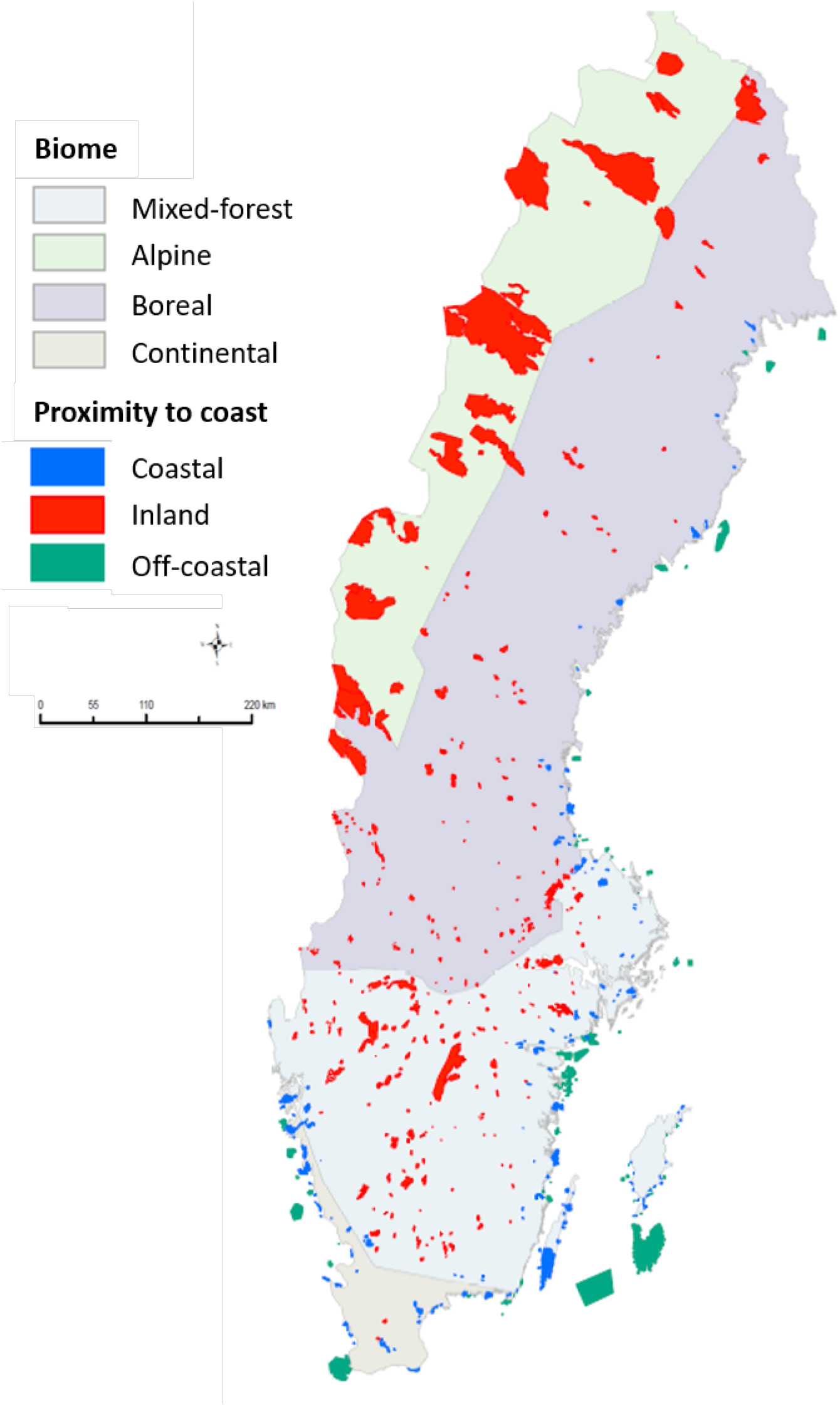
Special Protection Areas (SPAs) in Sweden divided by biogeographical region; see text for further explanation.

To assess the bird conservation value of SPAs that is lost due to infrastructure, I assumed an average of 20% (C.I. 12-33%) reduction of bird abundance within the 1 km effect distance from infrastructure, as indicated by the results presented by Benítez-López et al. (2010; see Fig. 1). GIS analyses were conducted using ArcGIS version 10.2 and QGIS version 2.6.

To explore how the degree of impact on an SPA is associated with its size and dominating habitat, two different analyses where performed for each region. For the probability of overlap with effect zone (response variable either 0 or 1), a generalized linear model with a logit link function (logistic regression) was used, and for the proportion of overlap with the effect zone (i.e., a response variable between 0 and 1), a beta-regression model (Ferrari & Cribari-Neto 2004) was used. In both types of models explanatory variables were 1) SPA size, 2) proportion forest habitat, 3) proportion wetlands, and 4) proportion agricultural land and grasslands. SPA size was log-transformed to improve normality and all variables were standardized to make parameter estimates comparable. Model selection was based on AIC and the final models were the ones with a combination of explanatory variables resulting in the lowest AIC. Statistical analyses were conducted using program R version 3.4.2.

## Results

The overlay of SPAs and effect zone of larger transport infrastructures showed that 339 of Sweden’s 538 SPAs (63%) have at least some part, and 123 (23%) have most of their area, within the effect zone (Table 1). In terms of area, a total of ca 126,000 ha, or 4.2% of the total SPA area in the country, lies within the effect zone.

National level figures on impacted area however gives a crude picture, as the results pointed at large differences among the biogeographical regions (Table 1). Alpine and marine areas have a number of large SPAs and hold most of Sweden’s total SPA area, but have sparse networks of large (terrestrial) transport infrastructures, and accordingly they reduce the national average. Continental, mixed-forest coastal and boreal coastal regions, however, are comparably more impacted; in these three regions combined a total of 46,046 ha, or 25.8% of the total SPA area, fall within the effect zone. In terms of bird conservation value, present infrastructure impacts are estimated to cause a 4-7% reduction in bird abundance in SPAs in the three most impacted regions (continental, mixed-forest coastal and boreal coastal), and an average of 1% reduction when all SPAs in the country are taken in consideration (Table 1).

The overlap models with the lowest AIC included SPA size or habitat composition in all regions except the continental (Table 2). As the most emergent pattern, larger SPAs appear to have a higher probability of overlap with effect zone (in three regions) but a lower proportion of overlap with effect zone (in all regions except continental). Another pattern, however less emergent, is that the overlap with effect zone is larger in SPAs with more agricultural land and grasslands (in four regions), and smaller in SPAs with more forest habitat (in two regions).

**Table 2.**
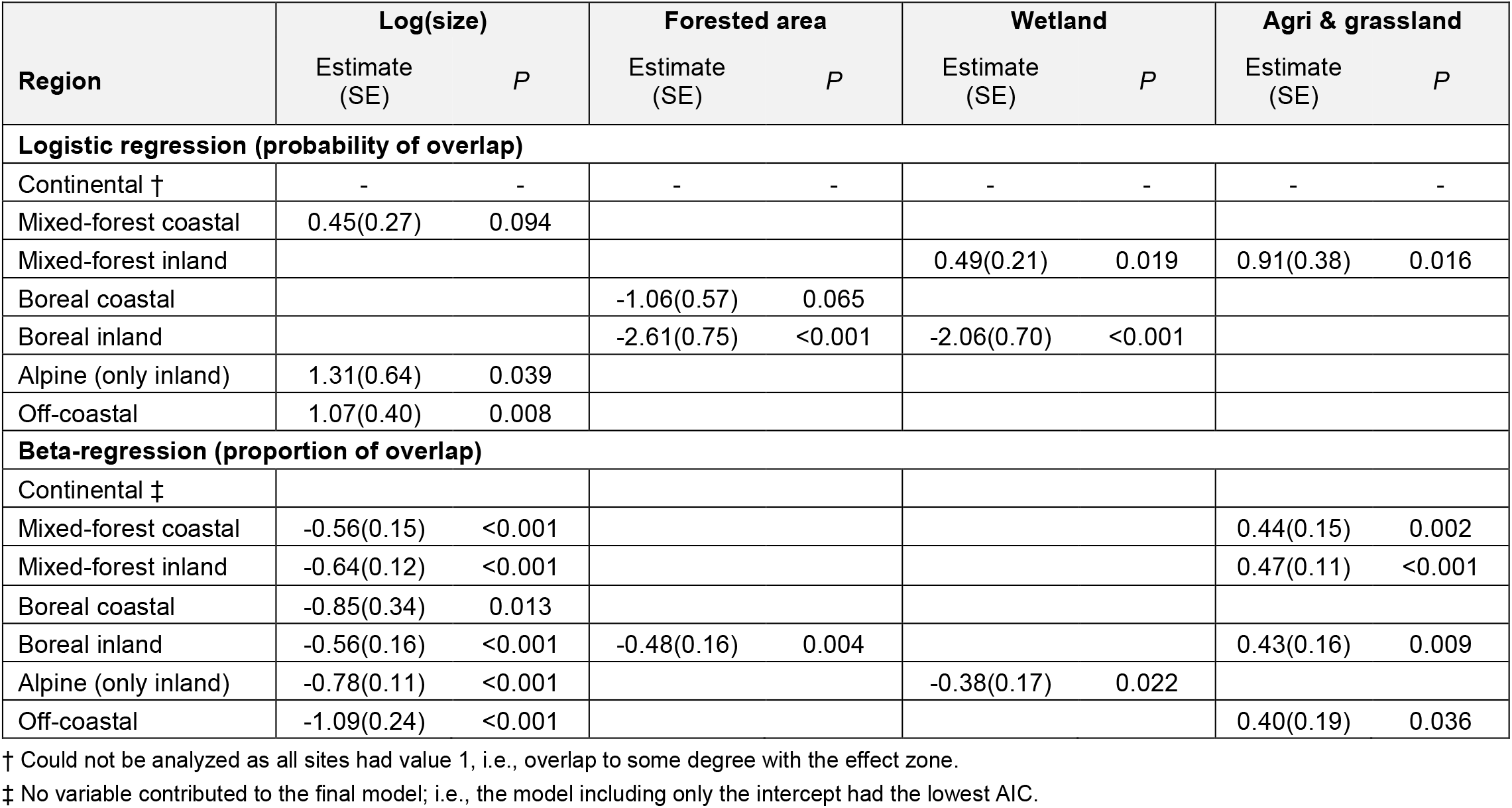
Variables associated with effect zone overlap with Swedish SPAs divided by biogeographical region. Values given are mean estimates of coefficients of logistic regressions and beta-regressions with standard errors (SE) and probabilities (P). Values are only given for variables that were included in the final model.

## Discussion

The results indicate that a significant proportion of Swedish SPAs, both in terms of area and number of sites, lies within a potential effect zone for birds around present larger transport infrastructures and therefore can be assumed not to reach their full potential as bird habitat. The reduction due to transport infrastructure impacts may not be dramatic when seen on the country as a whole, with only around 4% of the total SPA area affected, corresponding to a ca 1% reduction in predicted bird abundance within SPAs. However, for more urbanized parts of the country, with a denser infrastructure network, the predicted impact and reduction is nearly an order of magnitude larger and may well be one of the main factors determining bird abundance in protected areas. This is the case in the southern (continental) and coastal regions of the country, where the urbanization and landscape fragmentation is in level with that of most western and central European countries (EEA 2011).

At the level of individual SPAs, smaller sites tend to have a higher proportion of overlap with the effect zone and accordingly can be assumed to experience a greater impact than larger sites. This pattern emerge in all biogeographical regions, except the continental where most SPAs are small and indeed impacted to a large degree. In effect, the larger impact on smaller sites amplifies the bias against area protection in the lowlands, i.e., the southern and coastal regions. This is in line with general concerns previously expressed about the small size of many protected areas in Europe and about the impacts from transport infrastructure, traffic and other urban development in the landscape surrounding them (Shafer 1995, Gaston et al. 2008, Maiorano et al. 2008, Kati et al. 2015). However, as indicated by Helldin & Tytor (2017), these concerns are not well expressed in the management plans for Swedish SPAs, and appear particularly underestimated in the regions where the impacts are the largest.

The function of predicted bird abundance and distance to infrastructure described by Benítez-López et al. (2010) have previously been applied twice to assess the ecological impact of an infrastructure network on a larger geographical level. Karlson & Mörtberg (2015) presented an assessment of the impacts of roads on habitats of high diversity value in Sweden (irrespective of their protection status), concluding that natural grasslands and southern broadleaved forest are likely to be particularly impacted; on national level, 13-19% and 16-24% of the total areas of these habitats are found within predicted effect zone for birds. Torres et al. (2016) estimated a 19.0% (CI = 9.6–25.6%) reduction in national bird numbers due to transport infrastructure for Spain, with all land in view and not only protected areas. They too concluded that some habitats (most notably farmland and maritime wetlands) might be disproportionally affected by transport infrastructure. In relation to these previous assessments, the present study is unique in that it points out the impacts specifically on protected areas, i.e., areas where nature conservation should be a top priority.

The assessment was aimed to give a general picture, and was therefore simplified in several respects. A fixed-width effect zone is less realistic, as the actual effect depend on the local context, such as the habitat distribution, topography, species and ecological processes involved (Forman & Dubliner 2000, Ritter’s & Wickham 2003, Bilging & Duping-Giroux 2006) or the road characteristics (e.g., Reined & Fop pen 2006, Rytwinski & Fahrig 2015). Also the stronger reduction in bird densities near the infrastructure within the effect zone (Fig.

1) provides an opportunity to assess the decline in bird abundance within the effect zone in individual sites in greater detail than here conducted (e.g., Torres et al. 2016).

Furthermore, the analysis of impacts of different habitats within SPAs was rather coarse, since the Natura 2000 database does not provide habitat maps. Therefore, I could not explore to what degree EU priority habitats (habitat types of community interest; EEC 1992) are distributed disproportionally within the effect zone.

### Implications for management of protected areas

The present study underlines the concern about the impact of transport infrastructures on wildlife in protected areas in general (Forman & Deblinger 2000, Ament et al. 2008, Selva et al. 2011, Tsiafouli et al. 2013), and in the smaller areas in particular (Shafer 1995, Maiorano et al. 2008). Following article 4 of the Birds Directive, EU Member States must take appropriate steps to avoid habitat deterioration and significant species disturbance within SPAs (EEC 2010). Accordingly, a stronger emphasis on keeping natural areas free from the impacts of heavy traffic and new roads and railways has been proposed for conservation and transport planning (Selva et al. 2011, 2015, Laurance & Balmford 2013, IENE 2015).

Management plans for Natura 2000 sites should better acknowledge the threats to wildlife conservation caused by both present transport infrastructure and new development projects in and near sites (Cortina & Boggia 2014, Helldin & Tytor 2017). Assessments of the effectiveness of individual Natura 2000 sites in maintaining biodiversity should include monitoring of population density and demography of species of special conservation concern (Gaston et al. 2006). Managers of Natura 2000 sites have the opportunity to conduct more detailed assessments of predicted impacts from transport infrastructure based on habitat maps, species occurrences, and local road characteristics, to serve as a basis for priorities in conservation planning and action. Special attention to road effects should thus be paid in protected areas with a large overlap with the effect zone, in areas hosting particularly vulnerable taxa, and in areas with pronounced impacts on EU priority habitats.

Conservation authorities should secure that infrastructure owners and managers do their best to minimize the negative impact of nearby roads and railways and related traffic. Technical mitigation of impacts of transport infrastructure on birds could include preventing bird— vehicle collision (e.g., with flight diverters), planning the timing of infrastructure maintenance and construction work to avoid particularly sensitive periods, providing crossing structures (safe over-/underpasses), and reducing noise and visual impacts through walls, berms or adapted paving. Traffic calming, speed reduction or road closing (permanent or temporary) would also provide reductions in road mortality, disturbance and barrier effects. Finally, compensatory measures such as habitat improvement or additional area protection could reinforce remnant populations and restore vital ecological processes.

## Acknowledgements

I am grateful to Vadym Sokol for assisting in GIS data retrieval and GIS analyses, and to Victor Johansson for assisting in the statistical analyses. I am also grateful to Lars Nilsson for providing helpful comments on earlier drafts of this paper. I thank Ana Benítez-López and coworkers for kind permission to reproduce their published figure. The study was financed by the Swedish Transport Administration as part of the research program TRIEKOL (triekol.se).

